# SynBa: Improved estimation of drug combination synergies with uncertainty quantification

**DOI:** 10.1101/2023.01.24.524900

**Authors:** Haoting Zhang, Carl Henrik Ek, Magnus Rattray, Marta Milo

## Abstract

**Motivation:** There exists a range of different quantification frameworks to estimate the synergistic effect of drug combinations. The diversity and disagreement in estimates make it challenging to determine which combinations from a large drug screening should be proceeded with. Furthermore, the lack of accurate uncertainty quantification for those estimates precludes the choice of optimal drug combinations based on the most favourable synergistic effect.

**Results:** In this work, we propose SynBa, a flexible Bayesian approach to estimate the uncertainty of the synergistic efficacy and potency of drug combinations, so that actionable decisions can be derived from the model outputs. The actionability is enabled by incorporating the Hill equation into SynBa, so that the parameters representing the potency and the efficacy can be preserved. Existing knowledge may be conveniently inserted due to the flexibility of the prior, as shown by the empirical Beta prior defined for the normalised maximal inhibition. Through experiments on large combination screenings and comparison against benchmark methods, we show that SynBa provides improved accuracy of dose-response predictions and better-calibrated uncertainty estimation for the parameters and the predictions.

**Availability:** The code for SynBa is available at https://github.com/HaotingZhang1/SynBa. The datasets are publicly available (DOI of DREAM: 10.7303/syn4231880; DOI of the NCI-ALMANAC subset: 10.5281/zenodo.4135059).

## Introduction

With the increased use of small molecule drugs in monotherapy and drug combination treatments, off-target toxicity and resistance to treatments are becoming clear challenges. The use of combinations of drugs offers a possible solution to reduce toxicity minimising doses and bypassing resistance with alternative targeting. Thanks to the development of high-throughput approaches to screen drug-sensitivity in cell lines, dose-response data for a large number of combinations have been made available. Examples include the AstraZeneca-Sanger DREAM challenge (DREAM) (Menden et al. [2019]), NCI-ALMANAC (Holbeck et al. [2017]), DrugComb (Zagidullin et al. [2019]), and the screening data from the Wellcome Sanger Institute (Jaaks et al. [2022]). The availability of these datasets makes it possible to predict the effect of drug combinations from modelling drug-sensitivity data, leveraging information from biological features of the cell lines and chemical characteristics of the drugs (Bulusu et al. [2016]).

To understand how drugs can work synergistically when combined, models with a quantification framework are required. Traditional quantification frameworks usually involve a null surface, based on a number of set assumptions. The Bliss model is based on the Multiplicative Survival Principle (Bliss [1939]), whereas the Loewe model is based on the Dose Equivalence Principle (Loewe [1953]). In the past decade, parametric methods have started to emerge as alternatives to the above. These include MuSyC (Meyer et al. [2019], Wooten et al. [2021]), BRAID (Twarog et al. [2016]) and the Effective Dose model (Zimmer et al. [2016]). All these frameworks are based on different assumptions and parameterisations. As a result of this, the outputs from these models often disagree. Although there have been efforts to unify the frameworks, this is still an open problem for the field.

To define a common framework for drug combination, we need a less ambiguous definition of *synergy* (Tang et al. [2015]). *When a combination is said to be synergistic*, it is unclear whether it implies that the combination is desirable in terms of its *potency* or its *efficacy*. Potency is the amount of dosage required for a drug to produce a specified effect, whereas efficacy is the degree of the beneficial effect produced by the drug (Meyer et al. [2019]). A strong synergistic potency implies the toxicity may be reduced when the drugs are combined, which is crucial for avoiding overdose, whereas a strong synergistic efficacy implies that the combination increases the maximal possible effect. Both aspects are relevant for progressing with pre-clinical and subsequent clinical investigation. An ideal combination would be potent and effective. However, in clinical research there are situations where a drug partner is requested only for enhancement of potency and not to increase efficacy. Until recently, in quantification frameworks including Bliss, Loewe, BRAID and the Effective Dose Model, potency and efficacy are entangled within the concept of *synergy* as defined in the traditional quantification frameworks. To tackle this problem, Meyer et al. [2019] developed MuSyC, a framework that decouples potency and efficacy following the principles of the generalised Hill equation. Although the MuSyC approach is effective for modelling both potency and efficacy, the model does not fully explore the challenge of model-based estimation of uncertainty. There are various sources of uncertainty associated with drug-sensitivity modelling. Firstly, the biology of dose-response relationships is still unknown despite existing efforts. This leads to uncertainty associated with insufficient scientific knowledge. Secondly, there are systematic and random errors arising from the experimental procedures. Thirdly, biological variation exists among cell lines of the same disease, resulting in a further source of noise. Finally, uncertainty also stems from limited information that can be extracted due to the small size of available data (*epistemic uncertainty*).

Given these multiple sources of uncertainty, it is often impossible to reach an accurate estimate of the quantities of interest, e.g potency of a monotherapy, or the synergistic effect (in terms of either potency or efficacy) of a combination. Since the *best* single deterministic estimates might not be reached, here we focus on accurately quantifying uncertainty in the model as deviation from the true estimate given the noise (e.g. the credible range of model parameter estimates/predictions).

Most existing frameworks for drug combinations either do not compute the uncertainty or estimate it by standard error (Zimmer et al. [2016]) or parametric bootstrap (Wooten et al. [2021]), each of which contains unrealistic assumptions. The standard error is only accurate when a large number of samples is available, which is not the case for most pharmacology datasets. Similarly, for the parametric bootstrap, an accurate uncertainty estimation relies on a sufficiently large number of observations to be reasonably close to the truth. In either case, the uncertainty estimation will often be inaccurate due to the small data size. For this reason, here we approach uncertainty estimation with a Bayesian framework, which incorporates the uncertainty by treating all parameters of interest as probabilistic quantities. This enables us to continuously model the uncertainty in our estimates as the number of measurements grows, without becoming over-confident. Moreover, the estimated quantities of interest are obtained simultaneously with their uncertainties, making this approach computationally efficient.

In the current literature, there are examples that use a probabilistic model to incorporate uncertainty in their outputs. Hand-GP (Shapovalova et al. [2021]) is a non-parametric model based on the combination of the Hand model with Gaussian processes, providing more believable uncertainty estimation than MuSyC in some cases. However, Hand-GP does not incorporate the 1D Hill equation that imposed biological constraints useful for providing interpretable model outputs. For example, monotonicity of the monotherapy fitted curve is not enforced in Hand-GP, meaning it is possible that the model produces a dose-response surface that is unlikely to occur in the *in vitro* setting (Tansey et al. [2022]). More importantly, due to the non-parametric structure of Hand-GP, the parameters describing potency and efficacy are lost. The bayesynergy model (Rønneberg et al. [2021]) is a Bayesian framework that models synergistic interaction effects using Gaussian Processes, which provide uncertainty quantification. Although flexible, its formulation is based on the Bliss independence assumption that is biased against drug combinations with a moderate level of efficacy (Wooten et al. [2021]). The bayesynergy model also does not separate out synergistic potency and efficacy.

Here, we design a flexible Bayesian framework to infer synergistic effects of drug combination (**SynBa**) where (1) the classic Hill equation is preserved to produce estimates of efficacy and potency (2) the existing biological knowledge or insight from historical data may be conveniently added through the prior distribution over parameters in the model. In SynBa we will use MuSyC as a baseline framework to decouple synergistic potency and efficacy and add probabilistic inference to provide outputs and their associated uncertainty. This is to design a framework estimating the most favourable scores and efficiently provide optimal candidates to the drug discovery pipeline with actionable decision-making criteria derived from the model outputs. SynBa is also the first synergy framework that simultaneously provides principled uncertainty estimation, preserves the Hill equation and decouples the synergistic efficacy and potency.

With our approach, when a combination is predicted to be synergistic, a level of confidence will be quantified for this prediction, together with a level of improvement in efficacy or in potency. The associated uncertainty can guide decision on further laboratory experiments to proceed to the next phase. This will provide *actionable metrics* for the subsequent stages of the drug discovery pipeline, which is an unmet need.

## Methods

Our proposed method can be used for both analysing existing dose-response data and predicting unseen dose-response data for a given monotherapy *𝒟* = (*X, Y*) = {(*x*_*i*_, *y*_*i*_)} or a given combination *𝒟* = (*X, Y*) = {(**x**_*i*_, *y*_*i*_)}. The covariate can be a scalar *x*_*i*_ (in monotherapy) or a vector **x**_*i*_ (in combination) corresponding to the drug dosages. The response *y*_*i*_ can be defined as cell growth or inhibition of growth, depending on how the data are collected. In this study, we focus on inhibitory datasets, where a large dosage typically results in growth inhibition. In this case, *y*_*i*_ is defined as the percentage of growth-inhibited cells. Nevertheless, our method can be easily modified to accommodate the opposite setting where the drug response is enhancing growth with respect to the dosages.

Fig. 1 (A) illustrates a typical dose-response matrix for a combination from a screening, where the first row and column contain monotherapy data and the remaining entries contain combination data. The core aim of our method is to infer the synergistic potency and efficacy given such a matrix (or a vector in the case of monotherapy). To accomplish this, we designed **SynBa**, a **Ba**yesian framework for the inference of **Syn**ergistic effects of drug combinations. SynBa is defined by a prior distribution *p*(***θ***) for the parameters ***θ*** and a likelihood function *p*(*Y* | ***θ***, *X*) for the drug responses *Y*. The likelihood function describes the probability of the responses given the dosages *X* and the parameters ***θ***. The prior distribution encodes the existing belief or knowledge about the parameters. In monotherapy, we have ***θ*** = *{E*_0_, *E*_1_, *C, H, σ*}, whereas in combination, ***θ*** = {*E*_0_, *E*_1_, *E*_2_, *E*_3_, *C*_1_, *C*_2_, *H*_1_, *H*_2_, *α, σ*}. The likelihood function and parameters encode the shape of the dose-response curve (defined in Boxes 1 and 2 and described in more detail below).

**Fig. 1.**
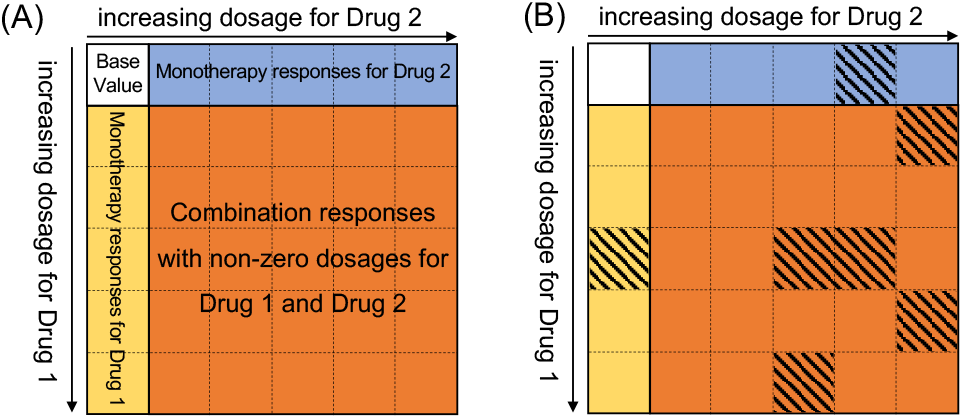
Description of the datasets for the proposed framework. (A): A typical dose-response matrix, where the top-left entry is the base value, the first row contains the monotherapy responses for the first agent, the first column contains the monotherapy responses for the second agent. The remaining entries are the responses when a combination of the two drugs is applied. (B): A dose-response matrix where not all responses are available for training. The shaded cells represent the test data.

In addition, our method provides a way of predicting unseen dose-response data for both monotherapies and combinations.

For example in the case of Fig. 1 (B), the quantity of interest would be the posterior predictive distribution of the response,

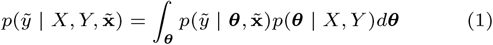

where 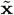 is the untested dosage of interest, 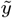 is the predicted response and *p*(***θ*** | *X, Y*) is the posterior distribution over the parameters ***θ*** given training data (*X, Y*).

## Overview of SynBa: Monotherapies

We begin by defining SynBa for monotherapy screens, where the dosage *x* is a scalar. The likelihood function for the response *y* is based on the Hill equation (Hill [1910]), which has been the classic choice to model pharmacology data:

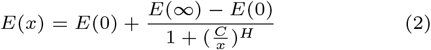

where *x* is the dosage of the drug, *E*(*x*) is the corresponding measured response, and *H* controls the slope of the curve. The interpretation of *C* depends on the problem and the dataset of interest. In this study, as the focus is on inhibitory datasets, *C* represents the dosage required to inhibit the given biological process or biological component by 50%, known as IC_50_. *C* quantifies the potency of the monotherapy in the study.

The base level of the monotherapy is denoted by *E*_0_ := *E*(0), which is the response when no drug is applied. The efficacy is quantified by *E*_1_ := *E*(*∞*), or denoted as E_inf_, which is the maximal inhibition when a sufficiently large dosage *x* is applied.

Box 1 defines the prior distribution for the parameters and the likelihood model for the response given the dosages, which are also illustrated in Fig. 2.

**Fig. 2.**
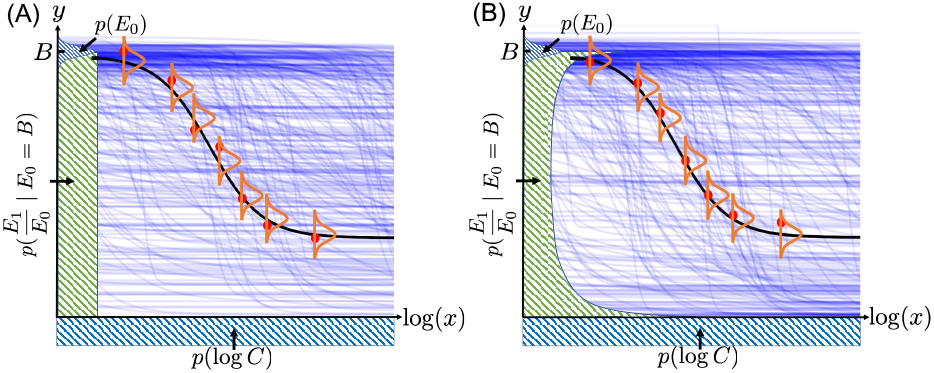
Two options (A) and (B) for the monotherapy prior model. The prior for log *C* is plotted underneath the x-axis, which is uniform in both options. The priors for *E*_0_ (which is Gaussian in both options) and the normalised *E*_1_ are plotted along the y-axis. Given *E*_0_, the normalised *E*_1_ is uniform in Option (A) and follows a Beta(0.46, 0.58) distribution in Option (B). The 300 blue curves are random samples from the expected prior responses E[*Y* |***θ***] where ***θ*** are sampled from the prior distributions defined in Box 1. The seven red points illustrate an example set of monotherapy dose-response data *D*. The black curve is a sample from the expected posterior responses E[*Y* |***θ***, *𝒟*] after the model is fitted to the data *𝒟*, whilst the orange bell-shaped curves illustrate the i.i.d. Gaussian noise for the responses.

To account for observational noise in the data, we define a noise model for *y* centred around *E*(*x*). For each fixed experiment setting (i.e. a fixed cell line treated by a fixed monotherapy or combination in the same laboratory environment), *y* is modelled as having Gaussian noise that is conditionally independent given the dosage: *y* ∼ *𝒩* (*E*(*x*), *σ*^2^). The assumption of conditional independence is valid because in screenings such as DREAM, the measurements for each monotherapy or combination are performed independently in different plates, instead of being performed sequentially. Therefore, given a set of measurements for a monotherapy or a combination, the noise level of their corresponding responses is independent. The i.i.d. noise is illustrated as orange bell-shaped curves in Fig. 2. As a result, the likelihood function for *y* is defined as Eq. (4).

The datasets are normalised by measuring cell inhibition when no drug is added (at dose zero). However, due to the noise in the biological process, the inhibition at dose zero would not be the same if the experiment is repeated multiple times. Thus, the normalization procedure itself contains uncertainty. Therefore, we define *E*_0_ to be probabilistic instead of a fixed initial value. A Gaussian prior with mean *B* and variance 0.03*B* is given for *E*_0_, where *B* is the normalized inhibition of a dose zero, e.g. *B* = 100 in DREAM. The variance of this prior is defined to be 0.03*B* so that it is flexible enough to allow for errors, but not too conservative.

The prior for the normalised maximal response (i.e. 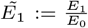, with a range of [0, 1]) may be defined in various ways depending on whether to insert existing knowledge or historical information. One option is to remain uninformative and impose a uniform prior, as shown in Fig. 2 (A).

Alternatively, we may make use of the existing information from the monotherapy data available. According to the single agent datasets in DREAM, the empirical distribution of E_inf_ has a high density on both extremes of the range, with 17.9% of them smaller than 0.05 and 15.1% equal to 1 (after normalising to the interval [0, 1]). Using maximum likelihood estimation to fit a Beta(*a, b*) distribution to these E_inf_ values, we obtain *a* = 0.46 and *b* = 0.58. To account for this information, we define Beta(0.46, 0.58) as the second option of the prior for the normalised maximal response, as illustrated in Fig. 2 (B). This prior is consistent with the biological behaviour that a drug administered with a sufficiently large dose will either kill the majority of the targeted cells if effective, or very few of them if not effective. The choice of this empirical prior shows how existing knowledge or information on the dynamic/kinetic of drugs can be conveniently added to SynBa through its priors.

A uniform prior is imposed for log *C* (the logarithm of IC_50_), with *C* bounded by *δ* and *M*. The values of *δ* and *M* depend on the dataset and the unit of the dosages. In this work, we define *δ* to be smaller than any non-zero dosage in the dataset

(i.e. 0 *< δ <* min{*x* | *x >* 0, (*x, y*) *∈ 𝒟*}) and *M* to be larger than any dosage in the dataset (i.e. *M >* max{*x* | (*x, y*) *∈ 𝒟*}). For example, *δ* = 10^*−*10^ and *M* = 10^6^ is a viable choice for DREAM, whereas in NCI-ALMANAC, we may have *δ* = 10^*−*15^ and *M* = 10. The idea is to be uninformative about log *C* a priori, since the ideal dosage range for the experiments is unknown and often unsuitable, either too small or too large. It is common that IC_50_ exceeds the maximum dosage. In other cases, IC_50_ may lie between zero dosage and the smallest non-zero dosage, due to the tested dosages being too large. Our method includes these possibilities a priori.

*H* and *σ* are both given a lognormal(0, 1) prior because they are both non-negative and assumed to be moderately small. In addition, previous literature indicated that *H* is approximately lognormal (Wooten et al. [2021]). A lognormal(0, 1) prior ensures that *P* (*H <* 5) *≈* 0.95 and *P* (*σ <* 5) *≈* 0.95 a priori.

The blue curves in Fig. 2 are 300 random samples from the expected prior responses 𝔼 [*Y* |***θ***] where ***θ*** are sampled from the prior distributions. It can be observed that the curves cover a wide range of possibilities a priori. Yet, they are not excessively general, as all of them follow the Hill equation.

Note that the priors we have chosen are motivated by general knowledge and previous literature. When more knowledge exists for a specific combination, the prior can be conveniently

### Box 1

Overview of SynBa for the inference of monotherapy dose-response data.

The joint prior distribution for the response and the parameters of the curve given the dosages is

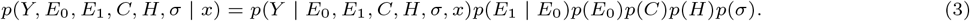

where the likelihood model for the responses *Y* given the dosage *x* is defined as

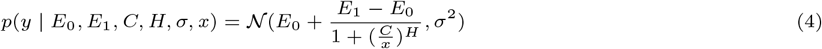

independently for all *y ∈ Y*, where *E*_0_ := *E*(0), *E*_1_ := *E*(*∞*) and *σ* is the standard deviation of the noise level of *y*, and the priors are

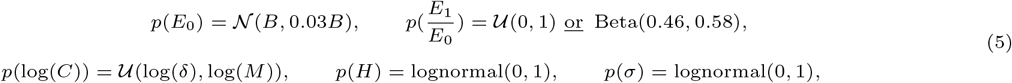

where 0 = *x*_1_ *< x*_2_ *≤ x*_3_ *≤* … *≤ x*_*n*_ are the dosages, and *δ ∈* (0, *x*_2_) is a small non-zero value to avoid log *C* being undefined. adjusted to accommodate this, since the inference framework is agnostic to the choice of prior.

Fig. 3 is an illustrative example of SynBa trained on a monotherapy (the compound MTOR 1 treated on the cell line MDA-MB-231) with six measurements, taken from the DREAM dataset. The first row shows the resulting model with a uniform prior for the normalised E_inf_ (i.e. 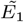), whereas the second row shows the model with the Beta(0.46, 0.58) prior for 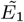. We start with two measurements and add two additional responses each time. It can be observed that the posterior distribution for IC_50_ narrows down quickly for both models because the observed responses span across the range between 40 and 100, which provides sufficient information to estimate IC_50_ with low uncertainty. The posterior for E_inf_, on the other hand, is more uncertain, due to the dosage range being too small to observe the convergence of the responses. In this case, the two models provide a visibly different posterior distribution for E_inf_, due to the different priors. As shown in Fig. 3 (F), the maximum a posteriori probability estimate for E_inf_ is 0 when the Beta prior is imposed, which results from the inserted prior knowledge that the response is likely to converge to 0 if an effective (but non-zero) response is already observed. This example shows how the prior design affects the inference of the parameter uncertainty. Nevertheless, if we look at the posterior predictive distribution for the responses (with samples illustrated by light blue curves), the two models reach a similar conclusion. We will show in Result that the predictive performance of the model is insensitive to the choices of prior.

**Fig. 3.**
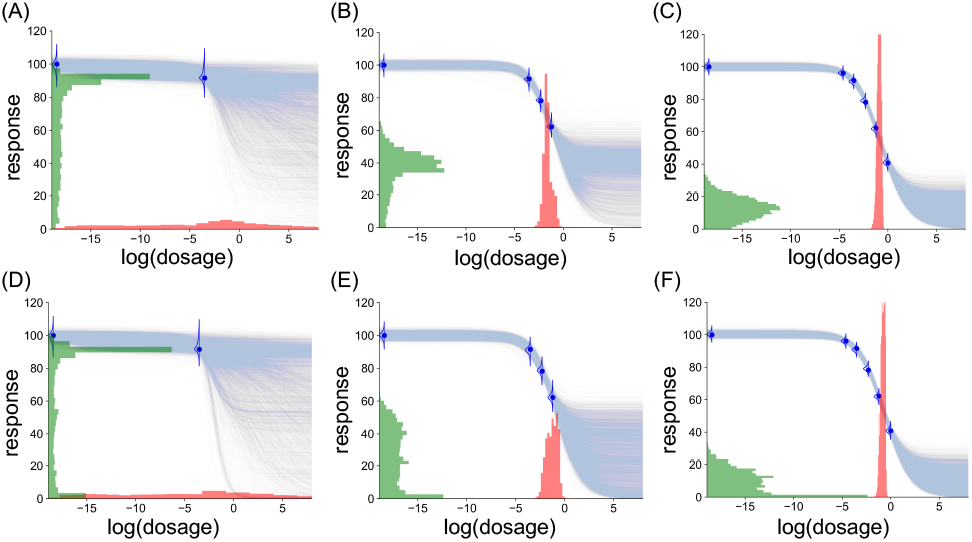
An example illustration of SynBa on a set of monotherapy data with six measurements. Each blue curve is a sample from E[*Y* | ***θ***] where ***θ*** *∼ p*(***θ*** | *𝒟*). The distribution for IC_50_ is shown in red, whereas the distribution for E_inf_ is shown in green. (A)-(C): The prior distribution for 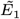 is *𝒰* (0, 1). (D)-(F): The prior distribution for 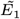 is Beta(0.46, 0.58).

## Overview of SynBa: Combinations

Extending SynBa to combinations of two drugs, the goal is now to model the dose-response surface *E*(*x*_1_, *x*_2_) = *f* (*x*_1_, *x*_2_) where *x*_1_ and *x*_2_ are the dosages for the two drugs, whereas *E*(*x*_1_, *x*_2_) is the response, and *f* is some class of function to be defined.

To define our likelihood model for the responses *Y*, we take inspiration from MuSyC but maintain some flexibility on their model assumptions. The effect of a drug in a system is usually described by the Hill equation that describes the state of equilibrium of a reversible process between an unaffected population and an affected one (the principle of *detailed balance*). To obey to this equation and its effects, in our model we incorporate the assumptions of the principle of detailed balance, of the proliferation rate of unaffected population and of the saturation of the maximum effect of the drug in the affected population. In MuSyC, these assumptions are defined in a nested structure in which levels are called *Tiers* (see Table S5 in Meyer et al. [2019]). In defining SynBa for combinations, we adopt the same model assumptions as *Tier 4* of the levels specified by MuSyC. This category encodes the most complex class of models that still maintains the assumption of *detailed balance*. It is worth noting that, conversely to MuSyC, our model posterior covers all four tiers simultaneously. This is because *Tier 4* subsumes all lower tiers and thus for how our model is defined, they will not be eliminated from the posterior distribution, unless the evidence from the data is strongly against them. The concepts of Tiers as described in MuSyC would require a post-learning model selection. The use of a Bayesian approach avoids such a selection procedure whilst still maintaining biologically viable assumptions.

In contrast to MuSyC, in SynBa we choose to maintain the detailed balance assumption to also avoid over-parameterisation, which is likely to occur due to the limited data size for each drug combination set. For example, the majority of the combinations in the NCI-ALMANAC dataset has a data size of 15 (excluding the base level at dose zero), only 3 more than the number of parameters in MuSyC. In a real-world scenario, the data size may often be even smaller. Furthermore, with the detailed balance assumption, matrix multiplication and inversion are avoided, which lowers the computational cost.

Box 2 defines the prior distribution for the parameters and the likelihood model for the response given the dosages.

### Box 2

Overview of SynBa for the inference of combination dose-response data.

We define the following joint distribution for the response and the parameters of the surface given the dosages **x** = (*x*_1_, *x*_2_),

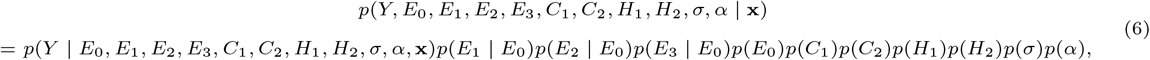

where the likelihood model for responses *Y* given the dosages is defined as

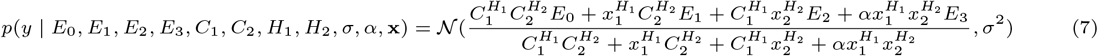

for all *y ∈ Y*, where *E*_0_ := *E*(0, 0), *E*_1_ := *E*(*∞*, 0), *E*_2_ := *E*(0, *∞*), *E*_3_ := *E*(*∞, ∞*), *C*_1_ and *H*_1_ are the monotherapeutic parameters associated for Drug 1, *C*_2_ and *H*_2_ are the monotherapeutic parameters associated for Drug 2, and *α* is an association parameter that controls how the two drugs are affected by the presence of each other.

The priors are

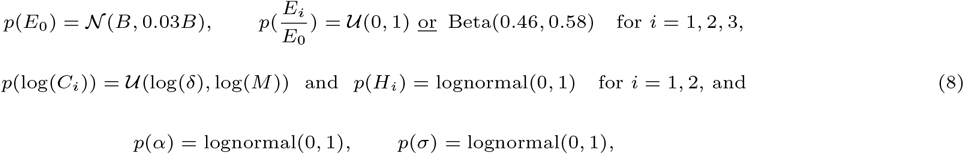

where 0 = *x*_*i*,1_ *≤ x*_*i*,2_ *≤* … *≤ x*_*i,n*_ are the dosages for drug *i*, and *δ* is a small non-zero value to avoid log *C* being undefined.

The definitions of the priors are a natural extension of the monotherapy model, with the same arguments being followed. The only new parameter is *α*, which follows a lognormal prior with median 1 because *α* is non-negative and equals 1 when the combination is neither synergistic nor antagonistic in terms of

### Inference of the synergy

After inferring the posterior for the parameters and their associated uncertainty, we focus on distinguishing the effect of efficacy and potency in drug combinations. MuSyC has defined metrics for both synergistic efficacy and synergistic potency, which is a promising step in decoupling potency and efficacy. However, the uncertainty for these two quantities has not been quantified systematically. Our model output includes not only quantification for the synergistic efficacy and the synergistic potency, but also a separate uncertainty estimation for each.

For the synergistic efficacy, one simple yet informative quantity is ΔHSA = min(*E*_1_, *E*_2_) *− E*_3_ which is the change in the maximal effect between the combination and the more effective single drug of the two (Greco [1995]). A positive score indicates synergistic efficacy. As *E*_1_, *E*_2_ and *E*_3_ are probabilistic, the resulting ΔHSA score is also probabilistic. A metric such as

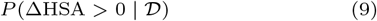

can then be defined to estimate how confident we are about the *synergistic efficacy* of the combination, based on the dataset *D*. It is possible to have a synergistic combination that is highly uncertain, which would indicate that more data are required to reach confidence in the estimation.

For the synergistic potency, *α* contains the required information. *α >* 1 would indicate synergistic potency (Wooten et al. [2021]), which means the potency of the two drugs has reduced due to being combined. Consequently, we define

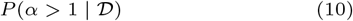

to estimate how likely a combination satisfies *synergistic potency*.

## Case studies

To illustrate how the uncertainty estimation from our method can be explained and further used for decision-making, we take two combinations from the DREAM dataset as examples.

Fig. 4 (A) shows the dose-response matrix for the combination of ADAM17 and AKT acted on the cell line BT-20. The first column is the monotherapy dose-response measurements for AKT (as the dosage for ADAM17 is zero), whilst the first row is the monotherapy measurements for ADAM17 (as the dosage for AKT is zero). It can be observed that the responses for AKT start to decrease at a higher rate when the dosage increases, but the dosage range is too small to understand its potency (IC_50_) and efficacy (E_inf_). The efficacy cannot be determined because the response has not shown any sign of convergence, whereas the potency cannot be determined because it relies on understanding the maximal response, which is itself uncertain. However, if a deterministic Hill equation is fitted to the monotherapy data, it will only provide a point estimate for the IC_50_ and E_inf_, without acknowledging the above caveats. On the contrary, our method provides an uncertainty estimation for both quantities. As shown in Fig. 4 (B)-(C), the posterior distribution of IC_50_ and E_inf_ have large variances, which correspond to large uncertainty. In particular, as shown in Fig. 4 (C), the posterior of E_inf_ for ADAM17 has a multimodal shape, which is sensible because it is unclear whether the dosage range is too small (which corresponds to the peak at 0), or the drug is simply ineffective regardless of the dosage (which corresponds to the peak at around 90).

**Fig. 4.**
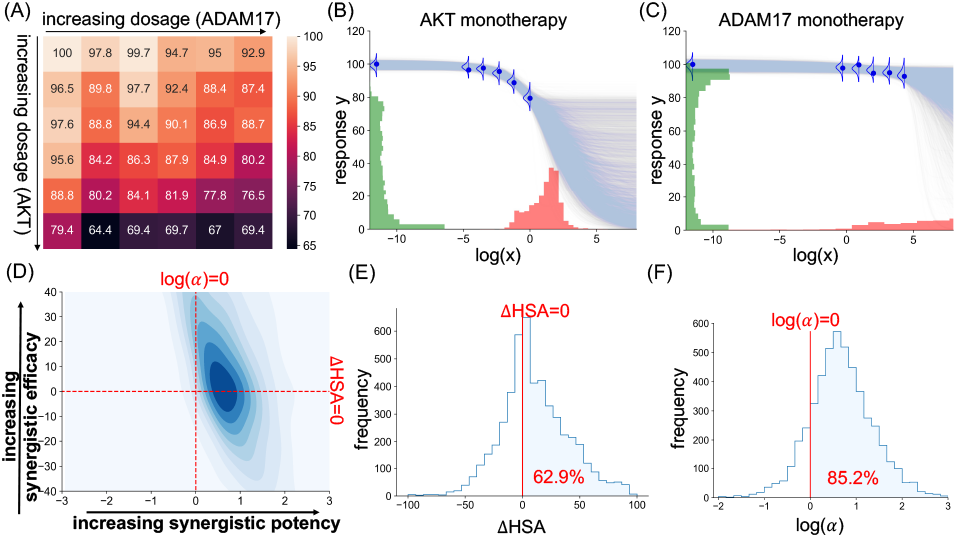
The data and the inference outputs of the combination of AKT and ADAM17 applied on the cell line BT-20. (A): The original dose-response matrix. (B)-(C): The monotherapy model outputs. Each blue curve is a sample from 𝔼 [*Y* | ***θ***] where ***θ*** is a sample from the posterior of the respective monotherapy model. The posterior distribution for IC_50_ and E_inf_ are shown in red and green respectively. (D): The contour plot for the joint posterior distribution of the synergistic efficacy (ΔHSA) and the synergistic potency (log(*α*)). The distribution is smoothed from the empirical posterior with a kernel density estimation for visualisation purpose. (E)-(F): The histogram of the empirical posterior distribution for the synergistic efficacy (ΔHSA) and the synergistic potency (log(*α*)). The areas on the right-hand side of the red vertical lines are the probability that the combination is synergistic in terms of efficacy (in (E)) and potency (in (F)).

Moving to the inference of the full combination matrix, most existing synergy methods have no means to showcase the uncertainty. Our method, on the contrary, provides the uncertainty around the synergistic potency and the synergistic efficacy, as shown in Fig. 4 (D), (E) and (F). According to the model output, the combination is moderately likely to be synergistically potent (with a probability of 85.2%), but it is difficult to conclude its synergistic efficacy (with a probability of 62.9% to be synergistic efficacious). This is reasonable because the excessively small dosage range makes it difficult to conclude anything about efficacy with high certainty, but with the 25 available measurements on the plates where the two drugs have interacted, information can be extracted on whether combining the two drugs may lower the level of toxicity required to reach the same beneficial effect.

This combination is an example where the model implies some potential in the synergy of the combination, but the level of uncertainty in the synergy is still high, which may require more measurements at larger dosages to be narrowed down.

We now consider the combination of AKT and EGFR acted on the cell line MDA-MB-468. Fig. 5 shows its dose-response matrix and the inference result from our model for the monotherapies and the combination respectively. All parameters and metrics of interest have low variances, representing low uncertainties. As shown in Fig. 5 (B) and (C), the dosages have suitable ranges and approximately follow the sigmoidal shapes of the expected dose-response fit, in particular for AKT. They contain sufficient information for the possibilities for IC_50_ and E_inf_ to be narrowed down. Similarly, the combination data are well-behaved. Fig. 5 (C), (D) and (E) show that the probabilities of this combination being synergistic in terms of potency and efficacy are both close to 100%. These are signs that this combination is worth being taken to subsequent steps in the drug development pipeline.

**Fig. 5.**
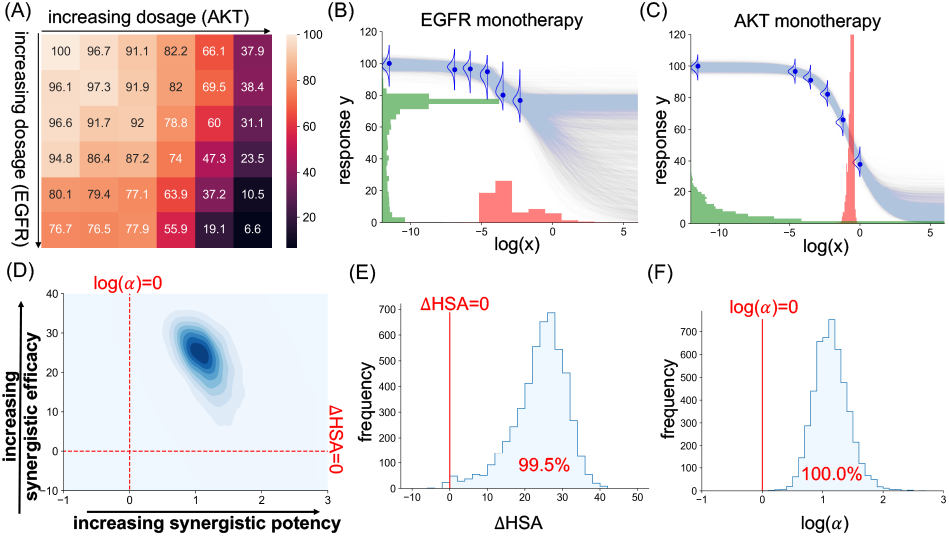
The data and the inference outputs of the combination of EGFR and AKT applied on the cell line MDA-MB-468. (A)-(F): The same as the caption of Fig. 4.

The two examples above show that concrete decisions can be made based on the posterior distributions (e.g. for IC_50_, E_inf_, ΔHSA and *α*) from our model, and more importantly, the uncertainties associated with these distributions.

## Training details

The models are trained by Stan, a state-of-the-art platform for statistical modelling and high-performance statistical computation, particularly for Bayesian computation (Stan Development Team [2023]). The user specifies the prior model of the parameters and the likelihood model of the data, while Stan performs either full Bayesian statistical inference with Markov chain Monte Carlo (MCMC) sampling, or approximate Bayesian inference with variational inference. In this study we opt for MCMC due to the importance of the reliability of the output, which is ensured by the asymptotic exactness of the MCMC inference. Despite choosing the slower option of the two, SynBa is still computationally efficient. Running on 4 CPUs of the Intel Xeon Platinum 8276 CPU Processor, the median time taken to fit SynBa (via MCMC with 1000 iterations and 4 chains, including 500 iterations in the warm-up phase) to a 6-by-6 dose-response matrix in DREAM is 10.2 seconds, which is comparable to MuSyC with bootstrap.

With this training pipeline, we can avoid the overhead that occurs during the usage of non-linear optimisation packages in deterministic parametric methods such as MuSyC, BRAID and the Effective Dose model. A different choice of the numerical algorithm (and its hyperparameters) results in a different result for those methods. On the contrary, in SynBa, the same exact result can be found asymptotically via MCMC with Stan.

For the implementation of the other benchmark methods including MuSyC, BRAID and the Effective Dose model, the Python package synergy (Wooten and Albert [2021]) is used.

## Results

### Prediction of drug combination responses

In this subsection, we show that in addition to providing uncertainty estimations, SynBa is competitive in predicting unseen responses within a dose-response matrix, and is less prone to overfitting compared to the existing methods.

The datasets of interest are DREAM (Menden et al. [2019]) and NCI-ALMANAC (Holbeck et al. [2017]), two of the most widely-used publicly-available combination screenings. In DREAM, we focus on all examples in the training set of Challenge 1 that have passed the Quality Assurance and that only contain non-negative responses and one set of replicates, which are 1631 sets of combinations in total. In NCI-ALMANAC, we focus on the subset defined in Julkunen et al. [2020], which is a subset of the data consisting of 50 unique FDA-approved drugs and 36,120 combinations. We remove examples that contain negative measurements, which results in 28,854 remaining combinations.

We leave out 20% of the non-zero dosage combinations for prediction. The models are trained using the remaining 80% dosage combinations, and then evaluated on the left-out points. For the dose-response matrices in DREAM, we leave out 7 of the 35 points with non-zero dosages from the 6-by-6 matrix (see Fig. 1(A)) using a specific leave-out strategy. For each of the two monotherapy slices, one point is left out for testing. For the 5-by-5 combination grid (i.e. the orange cells in Fig. 1(A)), five points are randomly left out for testing. Fig. 1(B) shows an example of such a train-test split. The measurements are left out in this manner so that each monotherapy contains one measurement for testing.

For the dose-response matrices in NCI-ALMANAC, it is not possible to leave out points separately for monotherapies and for combinations because the data size is too small. For most combinations, there are only 3 points for each monotherapy and 9 points for the interactions. Thus, we directly leave out 3 of the 15 points randomly for prediction.

### Evaluation metrics

To evaluate the predictive performance of the models, test likelihood and the root-mean-square error (RMSE) are computed using the left-out points. The former focuses on the goodness of the full predictive distribution, whereas the latter focuses solely on the goodness of the point estimates for the responses.

The computation of the test likelihood for SynBa follows Equation (1) for both monotherapy and combination. However, the right hand side of the equation cannot be computed in a closed form, so Monte Carlo estimation is required using the expression

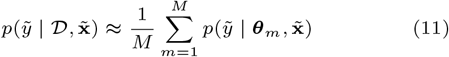

where 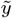 is the predicted response of the left-out dosage, *𝒟* is the training data containing the known dosages and the known responses, and ***θ***_*m*_ *∼ p*(***θ*** | *𝒰*) are the MCMC samples from the posterior distribution for the parameters.

For MuSyC, BRAID and the Effective Dose model, the parametric bootstrap pipeline described in Wooten et al. [2021] is followed. Each bootstrapped dataset provides a fitted curve. The density for 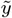 is then estimated by averaging its density computed on the models learnt from the bootstrapped datasets.

The computation of RMSE is more straightforward. For each combination, its RMSE for the test responses {*y*_1_, …, *y*_*N*_} is

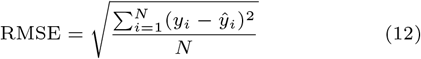

where *ŷ* _*i*_ is the point estimate for the response that corresponds to dosage **x**_*i*_. For SynBa, we define *ŷ* _*i*_ to be the posterior predictive mean 𝔼[*y*_*i*_ | *𝒟*, **x**_*i*_], which is estimated by Monte Carlo sampling from the posterior.

### Quantitative results

We compare our prediction results against MuSyC, BRAID, and the Effective Dose model, which are three of the most widely-used synergy models. For SynBa, we implement both the uniform prior and the empirical Beta prior for the normalised E_inf_, which we denote as SynBa-U and SynBa-B respectively. Tables 1 show the mean and the median of the test log-likelihood and the test RMSE for MuSyC, BRAID, Effective Dose model and SynBa. Our method outperforms all three other methods in all metrics except for the mean test log-likelihood on DREAM. In particular, our method performs well on RMSE, a metric that only considers the quality of point estimates and ignores uncertainty. The upper diagonal panels in Fig. 6 show the scatter plots directly comparing the test RMSE values between methods (visualised with the blue colour). Our method is the most competitive, as evidenced by having more points above the diagonal *y* = *x* line. These show that our method is not trading off predictive accuracy for uncertainty estimation. By following a principled Bayesian workflow, our model is strong in both prediction and uncertainty estimation.

**Table 1.** The mean and the median (Mdn) of the test log-likelihood (LL) and the test root-mean-squared error (RMSE) for MuSyC, BRAID, the Effective Dose model (ED) and SynBa, computed on a subset of DREAM and NCI-ALMANAC, along with their standard errors. The standard error of the mean is computed by the standard deviation of the metrics across examples divided by the square root of the number of examples. The standard error of the median is estimated by nonparametric bootstrap. SynBa with a uniform prior for the normalised E_inf_ is denoted by SynBa-U. SynBa with the Beta(0.46, 0.58) prior for the normalised E_inf_ is denoted by SynBa-B.

|  | DREAM, LL |  | DREAM, RMSE |  | NCI-ALMANAC, LL |  | NCI-ALMANAC, RMSE |  |
| --- | --- | --- | --- | --- | --- | --- | --- | --- |
| | mean (± se) | Mdn (± se) | mean (± se) | Mdn (± se) | mean (± se) | Mdn (± se) | mean (± se) | Mdn (± se) |
| MuSyC | $-3.50 \pm 0.35$ | $-3.09 \pm 0.02$ | $6.11 \pm 0.10$ | $5.01 \pm 0.08$ | $-3.86 \pm 0.01$ | $-3.79 \pm 0.01$ | $14.87 \pm 0.10$ | $10.17 \pm 0.03$ |
| BRAID | $-4.12 \pm 0.69$ | $-3.06 \pm 0.02$ | $5.71 \pm 0.09$ | $4.88 \pm 0.08$ | $-3.80 \pm 0.04$ | $-3.48 \pm 0.01$ | $9.57 \pm 0.06$ | $7.16 \pm 0.03$ |
| ED | **$-3.37 \pm 0.09$** | $-3.22 \pm 0.02$ | $6.46 \pm 0.09$ | $5.66 \pm 0.09$ | $-3.68 \pm 0.02$ | $-3.48 \pm 0.01$ | $8.47 \pm 0.05$ | $6.77 \pm 0.09$ |
| SynBa-U | $-3.59 \pm 0.42$ | **$-3.01 \pm 0.02$** | $5.20 \pm 0.07$ | $4.56 \pm 0.08$ | $-3.42 \pm 0.02$ | $-3.24 \pm 0.01$ | $6.72 \pm 0.04$ | $5.45 \pm 0.05$ |
| SynBa-B | $-3.84 \pm 0.66$ | **$-3.01 \pm 0.02$** | **$5.15 \pm 0.07$** | **$4.55 \pm 0.08$** | **$-3.39 \pm 0.01$** | **$-3.23 \pm 0.01$** | **$6.66 \pm 0.04$** | **$5.43 \pm 0.04$** |

**Fig. 6.**
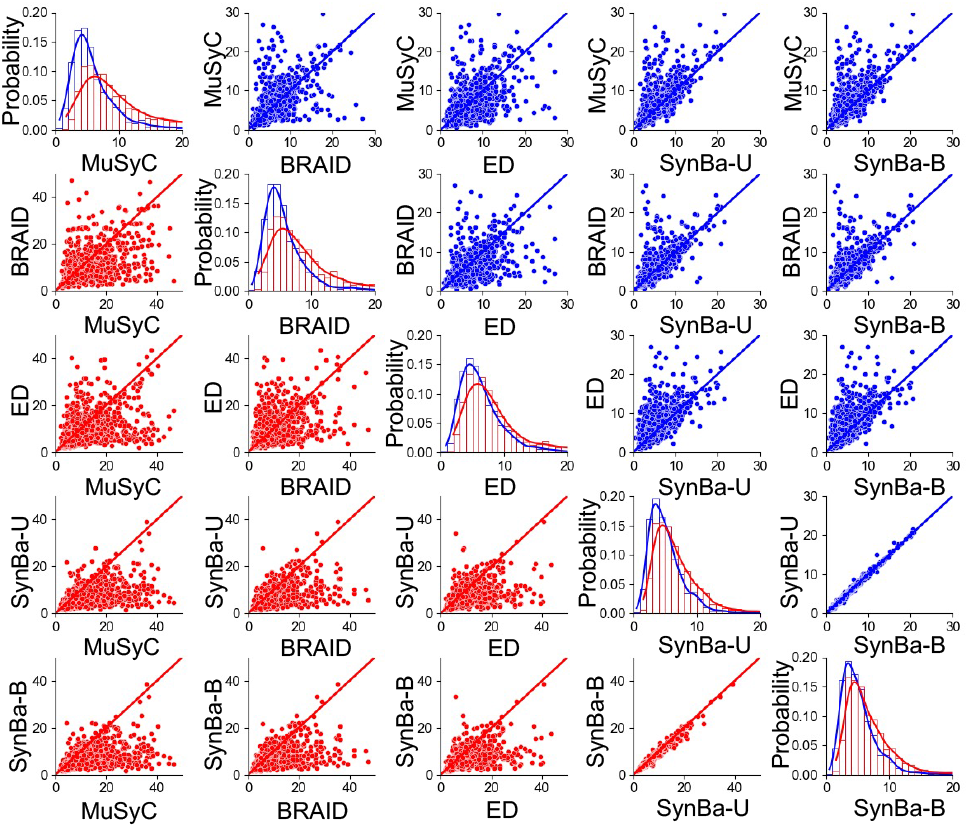
Upper diagonal panels (with blue points): Scatter plot of the test RMSE values obtained from different methods on DREAM with a train-test split ratio of 80% : 20%. Lower diagonal panels (with red points): Scatter plot of the test RMSE values obtained from different methods on DREAM with a train-test split ratio of 40% : 60%. Diagonal panels: Histograms of the test RMSE values and their corresponding kernel density estimates, where the train-test split is 4:1 for the blue ones and 2:3 for the red ones.

It is worth noting that at least one of MuSyC, BRAID or the Effective Dose model fail to find a solution for 4.8% of the examples in DREAM and 38.4% of them in NCI-ALMANAC, despite an effort in tuning the bounds, initial values and hyperparameters involved in the optimisation. Most likely this is because these methods rely on external optimisation packages with no guaranteed convergence, which can become a problem when overparameterisation becomes severe due to small data sizes. SynBa does not incur this problem since its priors ensure conservative outputs when data size is too small.

To investigate whether SynBa is prone to overfitting and how it compares to the other three methods, we perform the same prediction experiment on DREAM, but with a train-test split ratio of 40% : 60% instead. As shown in the lower diagonal panels in Fig. 6, the test RMSE values (visualised with the red colour) increase significantly for MuSyC, BRAID and the Effective Dose model. For SynBa, however, the test RMSE values have increased on average, but not by much. It can be seen that the mean value and the spread increase more significantly for the other three methods compared to SynBa. It can also be observed that the predictive performance of SynBa is not sensitive to the choice of prior, with the two priors producing very close RMSE values to each other.

In addition, we compare SynBa with bayesynergy and Hand-GP, the two probabilistic synergy models. As shown in Supplementary Table S1, the test RMSE for bayesynergy and Hand-GP are significantly higher on both datasets. This could be because drug interactions are modelled nonparametrically in both methods, meaning a wide range of functions is represented and it is difficult to narrow down the probable dose-response curves or surfaces when the data size is small or noisy.

## Uncertainty calibration

For a model *ℳ* with learnt cumulative distribution *F*_*ℳ*_ with well-calibrated uncertainty, it would approximately follow the identity that *F*_*Y*_ (**x**_*i*_) *≈ F*_*ℳ*_(**x**_*i*_) for every data point {(**x**_*i*_, *y*_*i*_(**x**_*i*_)) | *i* = 1, …, *N*} in the dataset, where *y*_*i*_(**x**_*i*_) is a sample from the unknown true cumulative distribution*F*_*Y*_ (**x**_*i*_). Equivalently, assuming the measurements *y*_*i*_(**x**_*i*_) for a combination are conditionally independent given the dosages **x**_*i*_, their cumulative probabilities *F* (*y*_*i*_) := ℙ(*y*_*i*_(**x**_*i*_) *< F*_*ℳ*_(**x**_*i*_)) would be approximately uniformly distributed between 0 and 1, if *ℳ* is well-calibrated.

In this study, for each combination in DREAM, we split the 35 measurements (excluding the base value) with a 80%:20% ratio in the same way as the prediction evaluation in the previous subsection. We then evaluate the quality of the uncertainty calibration with the Kolmogorov–Smirnov (K-S) uniformity test (Massey Jr [1951]) for the empirical cumulative probabilities (or CDF values) across all test data points. If the model is well-calibrated, then the CDF values for the test data points will be approximately uniform for each combination. Otherwise, they will show a non-uniform pattern and the resulting p-value for the K-S test will be statistically significant. This procedure is performed across every combination in DREAM, resulting in a p-value for each combination. The histogram of the p-values for SynBa is in Supplementary Fig. S1(A), showing that 6.07% of the combinations have not passed the uniformity test, and thus are not well-calibrated. As a comparison, the same procedure is performed using MuSyC. Supplementary Fig. S1(B) shows that 25.1% of the combinations are not well-calibrated when modelled by MuSyC, which is roughly four times as high as the number for SynBa. This shows the estimated uncertainty from SynBa is more reliable and closer to the unknown ground truth on average.

## Discussion

Machine learning methods have been developed for preclinical modelling and the prediction of drug combinations, thanks to the availability of large screenings (Julkunen et al. [2020], Wang et al. [2021]), which are beneficial for discovering and explaining drug combinations. However, a few factors prevent most from being applied to real-world drug discovery projects. One issue is that performance measures rely on synergy scores, which do not have a gold standard and contain a non-trivial amount of uncertainty, as discussed in Introduction. The Spearman correlation of the replicate experiments in DREAM (Menden et al. [2019]) and O’Neil et al. [2016] are 0.56 and 0.63 respectively, which show that quantifying a combination with a single synergy score would result in high variance. However, uncertainty measurement is not included in synergy score estimation. This could be one of the reasons that 20% of drug combinations are poorly predicted by all methods in the DREAM challenge. Measuring the uncertainties associated with the estimated scores is important for the subsequent decision-making process based on the model outputs. In real-world scenarios, scientists are often facing the decision to choose amongst a large set of drug combinations that score similarly in terms of synergy. Without quantification of how certain (or uncertain) the estimated scores are, they will have to rely on background knowledge compromising innovation in their choices. SynBa provides a way to implement a ranking strategy in the decision process of a drug discovery pipeline, which is a real-world unmet need.

SynBa has the limitation that it only models the combination of two drugs, while there exist methods such as the Effective Dose Model that consider higher-order combinations. While this is a limitation, our initial aim is to provide a method that is reliable and not over-parameterised, to meet with practical needs.

## Conclusions

We have developed a new framework for quantifying dose-response relationships for monotherapies and combinations that provides a full uncertainty estimation for all parameters that are associated with the monotherapies and the combinations, including information about efficacy, potency and synergy. These uncertainty information would be helpful to the biologists to make further decisions about progressing to the next stages of the drug discovery pipeline, or whether more experiments are required to lower the level of the uncertainty and better understand the drug mechanism of action.

We have also shown that SynBa is competitive in predicting unseen responses within a given dose-response matrix, and outperforms MuSyC, BRAID and the Effective Dose model on DREAM and NCI-ALMANAC. In addition, the prediction performance is not sensitive to the choice of the priors.

In summary, our framework is capable of providing a reliable uncertainty estimation for the potency (e.g. IC_50_) and the efficacy (e.g. E_inf_) of a monotherapy, or the synergistic potency and efficacy of a combination, in a decoupled manner, and reliably predicting unseen responses within a dose-response matrix. The parameter uncertainties can be interpreted and used as guidance for further experiments and subsequent decision-making.

## Acknowledgments

The authors thank Mateja Jamnik, Pietro Liò, Krishna Bulusu, Paul Metcalfe and Elizaveta Semenova for their valuable comments and suggestions. This work was performed using resources provided by the CSD3 operated by the University of Cambridge Research Computing Service.

