## Supplementary Materials (Table S1 and Figure S1) for "SynBa: Improved estimation of drug combination synergies with uncertainty quantification"

March 23, 2023

### 1 Supplementary Figure

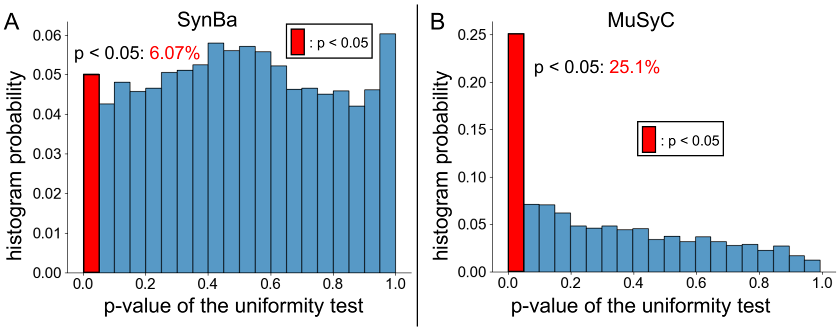


Fig. S1: Histogram of p-values that represent the calibration quality of combination models. The p-values are derived from the Kolmogorov–Smirnov test between the uniform distribution and the cumulative probability of the data points in their predicted densities. For a well-calibrated model, the p-value will be higher than 0.05.

### 2 Supplementary Table

Table S1: The mean and the median of the test root-mean-squared error (RMSE) for MuSyC, BRAID, the Effective Dose model, bayesynergy, Hand-GP and SynBa, computed on a subset of DREAM and NCI-ALMANAC, along with their standard errors. The standard error of the mean is computed by the standard deviation of the metrics across examples divided by the square root of the number of examples. The standard error of the median is estimated by nonparametric bootstrap. SynBa with a uniform prior for the normalised $E_{inf}$ is denoted by SynBa-U. SynBa with the $Beta(0.46, 0.58)$ prior for the normalised $E_{inf}$ is denoted by SynBa-B.

| | DREAM, RMSE | | NCI-ALMANAC, RMSE | |
|---|---|---|---|---|
| | mean ($\pm$ se) | median ($\pm$ se) | mean ($\pm$ se) | median ($\pm$ se) |
| MuSyC | $6.11 \pm 0.10$ | $5.01 \pm 0.08$ | $14.87 \pm 0.10$ | $10.17 \pm 0.03$ |
| BRAID | $5.71 \pm 0.09$ | $4.88 \pm 0.08$ | $9.57 \pm 0.06$ | $7.16 \pm 0.03$ |
| Effective Dose | $6.46 \pm 0.09$ | $5.66 \pm 0.09$ | $8.47 \pm 0.05$ | $6.77 \pm 0.09$ |
| bayesynergy | $8.10 \pm 0.03$ | $8.07 \pm 0.04$ | $24.18 \pm 0.12$ | $17.92 \pm 0.20$ |
| Hand-GP | $10.02 \pm 0.16$ | $8.35 \pm 0.12$ | $22.91 \pm 0.31$ | $14.41 \pm 0.37$ |
| SynBa-U | $5.20 \pm 0.07$ | $4.56 \pm 0.08$ | $6.72 \pm 0.04$ | $5.45 \pm 0.05$ |
| SynBa-B | $\mathbf{5.15} \pm 0.07$ | $\mathbf{4.55} \pm 0.08$ | $\mathbf{6.66} \pm 0.04$ | $\mathbf{5.43} \pm 0.04$ |
